# A transcription factor and a phosphatase regulate temperature-dependent morphogenesis in a fungal plant pathogen

**DOI:** 10.1101/2022.02.07.479299

**Authors:** Carolina Sardinha Francisco, Bruce A. McDonald, Javier Palma-Guerrero

## Abstract

Naturally fluctuating temperatures provide a constant environmental stress that requires adaptation. Some fungal pathogens respond to heat stress by producing new morphotypes that maximize their overall fitness. The fungal wheat pathogen *Z. tritici* responds to heat stress by switching from its yeast-like blastospore form to hyphae or chlamydospores. The regulatory mechanisms underlying this switch are unknown. Here, we demonstrate that a differential heat stress response is ubiquitous in *Z. tritici* populations around the world. We used QTL mapping to identify a single locus associated with the temperature-dependent morphogenesis and we found two genes, the transcription factor *ZtMsr1* and the protein phosphatase *ZtYvh1*, regulating this mechanism. We find that *ZtMsr1* regulates repression of hyphal growth and induces chlamydospore formation whereas *ZtYvh1* is required for hyphal growth. We next pinpointed that chlamydospore formation is a response to the intracellular osmotic stress generated by the heat stress. This intracellular stress stimulates the CWI and HOG MAPK pathways resulting in hyphal growth. If cell wall integrity is however compromised, *ZtMsr1* represses the hyphal development program and might induce the chlamydospore-inducing genes as a stress-response survival strategy. Taken together, these results suggest a novel mechanism through which morphological transitions are orchestrated in *Z. tritici* – a mechanism possibly also present in other pleomorphic fungi.

**IMPORTANCE:** Temperature is an environmental signal constantly monitored by pleomorphic fungi. Our experiments showed that yeast-to-hyphal or yeast-to-chlamydospore transitions are ubiquitous heat stress responses in *Z. tritici*. QTL mapping allowed us to identify a transcription factor and a protein phosphatase contributing to temperature-dependent morphogenesis. We showed that intracellular osmolarity is the pivotal signal inducing these transitions. We propose a regulatory network controlling *Z. tritici* morphogenesis, which may have broad implications for temperature sensing of fungal pathogens.

## INTRODUCTION

Temperature fluctuation is a ubiquitous stress that disrupts homeostasis in all organisms, including fungi. Abrupt temperature change demands a rapid re-adjustment of fungal physiology and therefore represent an environmental stress signal. Some fungal species respond to heat stress by altering their growth forms. For these fungi temperature shifts provide decisive environmental cues that affect their development. For example, thermally dimorphic human pathogens, such as *Histoplasma capsulatum* and *Paracoccidioides brasiliensis*, grow as hyphae at ambient temperatures below 30°C and convert into the pathogenic yeast form at the elevated host temperature [1-3]. In contrast, the ambient temperature favors the yeast-like phase of *Candida albicans*, while high temperature induces filamentous growth [4]. Though the morphological outcomes seem to be tightly associated with the life histories of different fungal species, the regulatory circuits controlling heat stress responses usually include the upregulation of specific thermal proteins (e.g., heat shock proteins - HSPs) [5, 6]; activation of mitogen-activated protein kinase (MAPK) signaling pathways (e.g., cell wall integrity (CWI) and high-osmolarity glycerol (HOG) pathways) [7-9]; and changes in cellular composition (e.g., glycerol production, trehalose accumulation, and increased chitin synthesis) [10-13], consistent with the idea that fungal cells have evolved shared mechanisms to cope with changing temperatures [14-16].

MAPK signal transduction pathways link environmental changes to transcriptional regulation in many eukaryotic cells. Most filamentous fungi use five different MAPK pathways in response to environmental signals [17]. Two of them, the CWI and HOG pathways, are regulated in a coordinated manner during heat stress [8, 10, 18, 19]. CWI is activated by perturbations of the cell surface or plasma membrane and is responsible for maintaining cell morphology [7]. Cell sensors at the fungal plasma membrane convey cell surface signals to the nucleus through sequential phosphorylation of MAPK proteins, which in turn promote the activation of MAPK *Slt2* (homologue *Mpk1* in *Saccharomyces cerevisiae*) [7, 20, 21]. Null mutants in the CWI pathway display altered growth, cell lysis defects, and thermosensitivity [22-28]. In its turn, HOG is a well-known regulatory pathway involved in responses to osmotic stress [29]. Osmoregulation maintains an appropriate intracellular environment for biochemical reactions and fungal cell turgor by controlling the efflux and influx of osmolytes [30]. Nevertheless, the HOG pathway is also involved in morphogenesis and cell wall biogenesis [29], two essential traits related to fungal virulence [31]. Thus, to define the cellular machinery that controls morphological transitions at high temperatures can provide unique insights into fungal stress responses and morphological adaptations.

The ascomycete fungus *Zymoseptororia tritici* was the first plant pathogen shown to undergo morphological transitions in response to high temperatures [32, 33]. *Z. tritici* is distributed globally and is the most damaging fungus in European wheat crops [34]. It exhibits a striking morphological variation, whereas *Z. tritici* strains morphologically respond differently to the same environmental conditions [35]. For example, temperatures ranging from 15°C to 18°C induce blastosporulation (blastospore replication by budding – generating a “yeast-like form”) while elevated temperatures (from 25°C to 28°C) promote transition to hyphal growth, pseudohyphae or chlamydospore production [33, 35]. Pseudohyphae are distinguished from true hyphae by constrictions formed at the septal junctions, leading to a loss in cytoplasmic continuity between cells [36]. In turn, chlamydospores are spherical thick-walled cells, and chlamydospores produced by *Z. tritici* are better able to survive several stresses that kill other cell types [35]. All these morphotypes can be produced during the asexual vegetative growth phase of the fungus [35], but it is unclear how the transition among these modes of growth are regulated. The natural morphological variability observed in pleomorphic fungi is often explained by variation in the genetic background, usually by a combination of a particular set of alleles that contribute to the phenotype in question. Given the complexity of this trait and the natural variation in morphological responses found within field populations of *Z. tritici* [35], we hypothesized that alleles associated with temperature-related morphotypes can be identified using quantitative trait locus (QTL) mapping.

Here, we report the morphological variability of five worldwide field populations of *Z. tritici* in response to heat stress. We conducted a QTL mapping study using a mapping population derived from parental strains exhibiting contrasting morphological responses. We identified a genomic region responsible for temperature-dependent morphogenesis in *Z. tritici*, and we functionally characterized the genes within this region. Among them, we found that the homolog of Yvh1, a well-known protein phosphatase and here termed *ZtYvh1*, is required for hyphal growth. We also found a transcription factor termed *ZtMsr1* functioning as hyphal repressor and chlamydospore inductor. Though both genes contribute to morphological transitions in *Z. tritici*, the *ZtMsr1* plays a major role in the differential heat stress response observed between the parental strains. Finally, we discovered that chlamydospore formation is a response to the intracellular osmotic stress generated during heat stress rather than being a response to the temperature upshift itself. Our experiments elucidate a novel regulatory circuit controlling differential morphotype transitions in a fungal plant pathogen, and illuminate the complex genetic architecture underlying temperature-dependent morphogenesis in pleomorphic fungi.

## RESULTS

### The global distribution and genetic architecture of temperature-dependent morphotypes

We previously reported that two field isolates of *Z. tritici* differ in their production of chlamydospores under heat stress [35]. Here, we expand upon this finding by assessing the morphological responses of 141 *Z. tritici* isolates from five worldwide field populations under heat stress (27°C) (see Table S1 and Fig. 1). In four out of five populations the majority of the isolates exhibited hyphal growth, except for the Israeli population, where 77% (n=23) of the isolates switched to chlamydospores at high temperatures (Fig. 1A). Regardless of geographical origin, chlamydospore- or hyphal-forming isolates shared the same morphological features (Figs. 1B-C), including the spherical and thickened cell walls characteristic of chlamydospores and the elongated cell walls lacking constrictions in hyphae [37].

**FIG 1.**
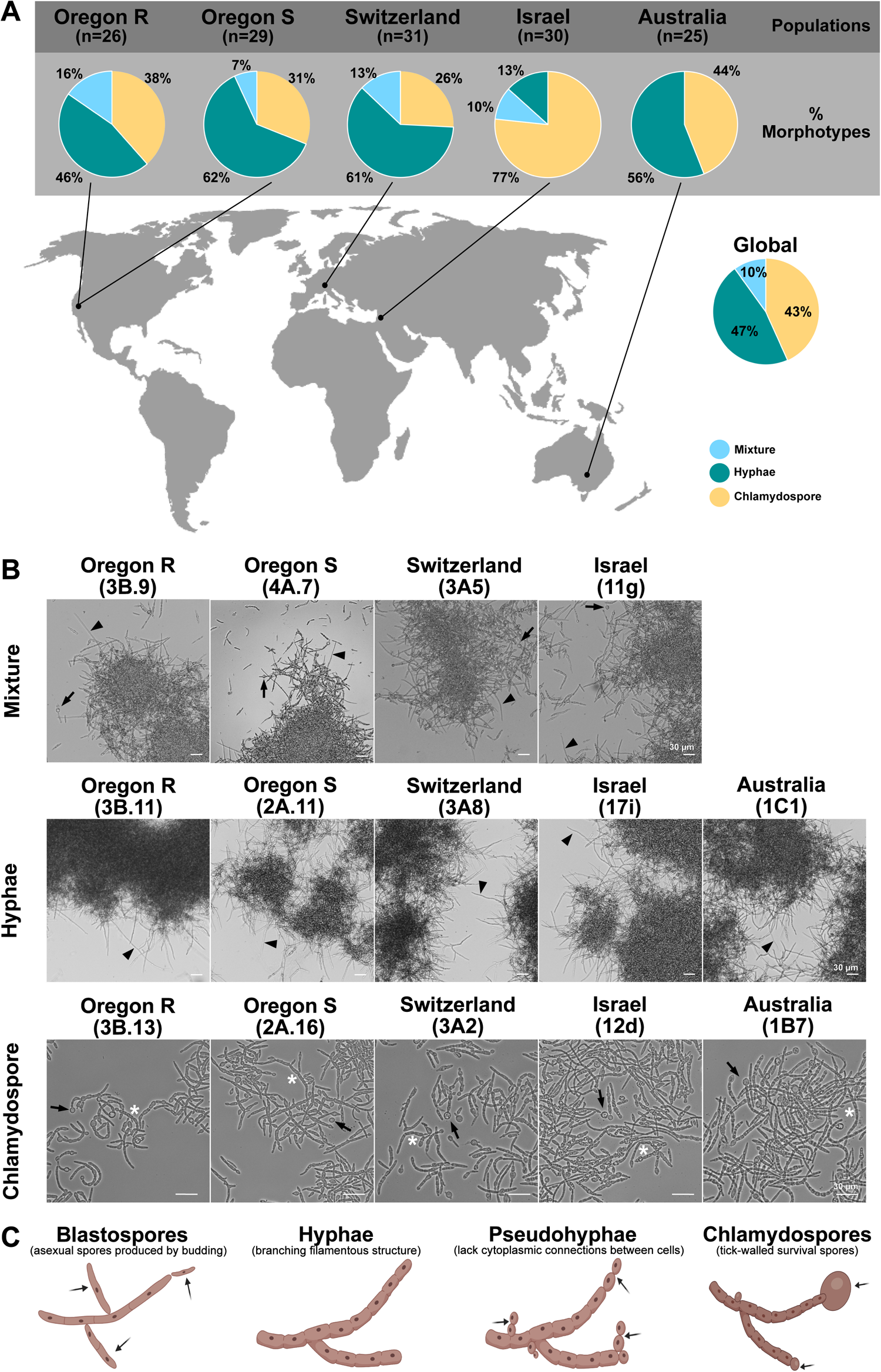
Global field populations of *Zymoseptoria tritici* vary in their heat stress response. (A) Sampling locations and associated frequency of isolates growing as chlamydospores or hyphae upon exposure to elevated temperature (27°C). Individuals that exhibited both morphotypes at high temperatures were termed mixture. (B) Micrographs of typical morphotypes of mixture- (upper panel), hyphal- (middle panel), or chlamydospore-growing isolates (lower panel) of each population. Black triangles indicate hyphae growth, white asterisks point to pseudohyphae, and black arrows demonstrate the chlamydospore cells. (C) Illustrations of the four morphotypes produced during vegetative growth of *Z. tritici*, such as blastospores, hyphae, pseudohyphae, and chlamydospores. Image created with BioRender.com.

To analyze the genetic architecture of temperature-dependent morphogenesis, the segregating F1 population from a cross between 1A5 and 1E4 was scored for its morphological response to grow as hyphae, chlamydospores, or both morphotypes at 27°C. The observed morphotype for each offspring is given in Table S2 and summarized in Fig. 2B. A genome scan of the F1 population identified a single QTL on chromosome 12 with a logarithm of odds (LOD) score of 17.8 (Figs. 2A and C). The QTL confidence interval was determined by the pseudomarkers at chr12.loc322 (LOD = 17.3), chr12.loc325 (LOD = 17.8), and chr12.loc329 (LOD = 16.8), which were flanked by the true SNP markers 12_1226988 (chr12.loc316) on the left and 12_1258185 (chr12.loc334) on the right side of the QTL (Fig. 2D). A detailed screening of the chromosome 12 QTL identified a 95% confidence interval of 31 Kb containing only eight genes (Fig. 2E and Table S3).

**FIG 2.**
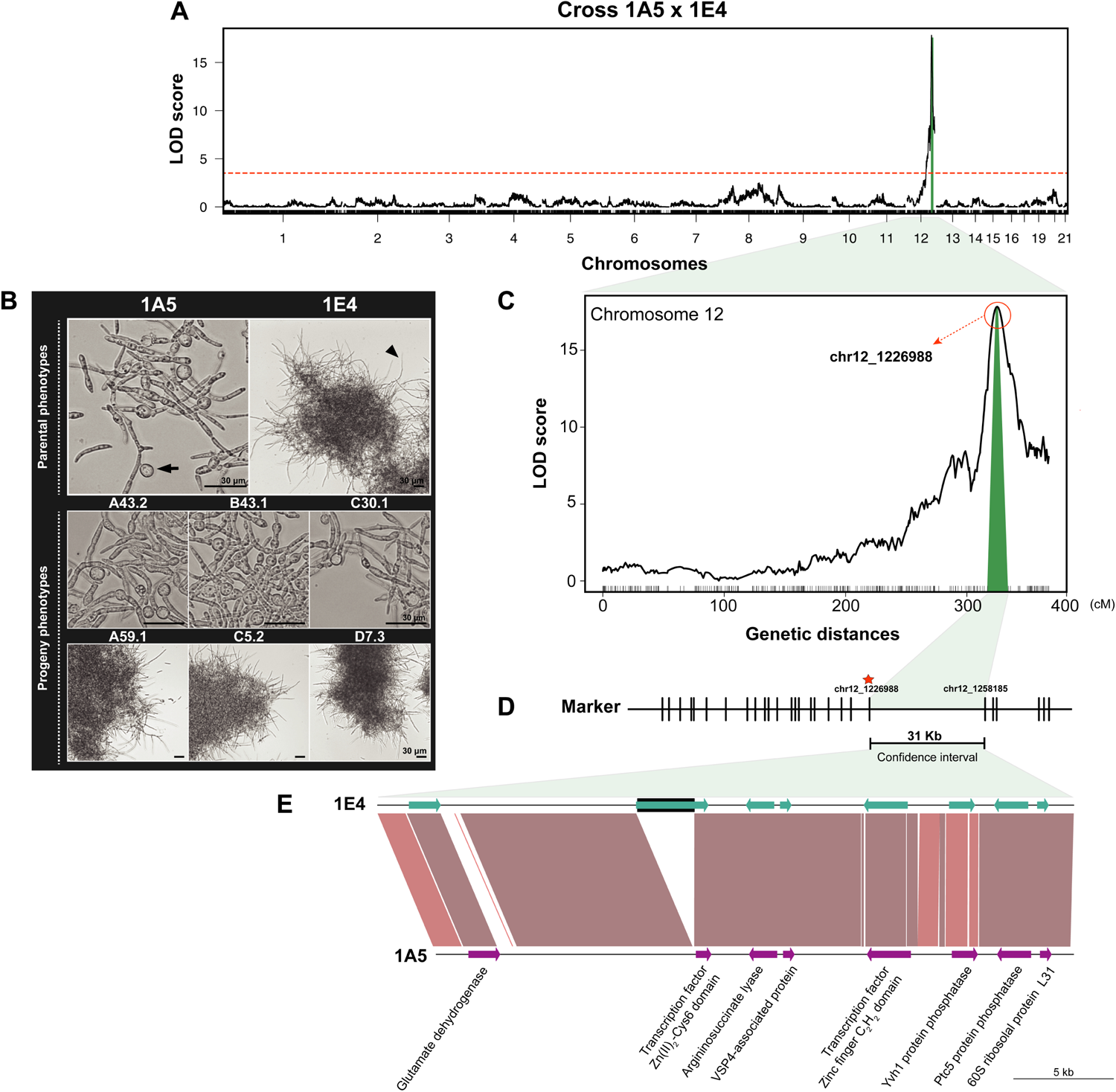
A QTL on chromosome 12 is correlated to temperature-dependent morphogenesis. (A) Interval mapping for the 1A5 (morphotype chlamydospore) x 1E4 (morphotype hyphal) cross. The y-axis shows the log_10_ of the odds (LOD score) values, and the x-axis indicates the chromosome numbers. The dashed horizontal red line represents the significance threshold (*p* = 0.05). (B) Representation of the morphological responses displayed by the parental strains and individuals from the segregating F1 population. The 1A5 parental strain produces chlamydospores (black arrow) without any evidence of hyphal differentiation, while 1E4 undergoes mainly filamentation in response to heat stress. (C) LOD plot of the chromosome 12 QTL. The x-axis indicates the genetic distances (centimorgans - cM) of the chromosome 12, including the chromosome 12 QTL 95% confidence interval (green color). (D) SNP markers surrounding the confidence interval are shown as vertical lines. The 12_1226988 marker on the left and the 12_1258185 marker on the right delimited the 31 kilobase (Kb) confidence interval size of the QTL. A red star indicates the true SNP marker (12_1226988) closest to the pseudomarker chr12.loc325 (LOD = 17.83) with the highest LOD score. (E) Synteny plot showing the genes located within the chromosome 12 QTL and DNA polymorphism in this genomic region. The darker the red color, the lower the degree of polymorphism between 1A5 and 1E4 parental strains. The arrows represent the genes and the black square indicates a transposable element.

### Four candidate genes were identified within the QTL confidence interval

Among these genes, we focused on those predicted to encode a gene function related to cell development and containing at least one sequence variant either in the protein-encoding sequence or the 5’ or 3’ UTRs. The four genes meeting these criteria were *1A5*.*g11034, 1A5*.*g11037, 1A5*.*g11038*, and *1A5*.*g11039* (Table S3). Two of the genes encoded putative transcription factors (TFs) that we named *ZtMsr1* (*<u>m</u>orphological <u>s</u>tress response 1*) (*1A5*.*g11034*) and *Zt11037* (*1A5*.*g11037). ZtMsr1* and *Zt11037* encode proteins containing GAL4-like Zn(II)_2_Cys6 and C2H2 zinc finger domains, respectively, but orthologs have not yet been characterized. The other two genes encoded protein phosphatases named *ZtYvh1* (*1A5*.*g11038*) and *ZtPtc5* (*1A5*.*g11039*). *ZtYvh1* is orthologous to the dual-specificity protein phosphatase Yvh1 that was first identified as a yeast vaccinia virus VH1 phosphatase in *Saccharomyces cerevisiae* [38], and has since been characterized in other fungi [39-41]. ZtPtc5 is orthologous to Ptc5 of *S. cerevisiae*, a protein located in the mitochondrial compartment [42, 43]. *Zt11037, ZtYvh1*, and *ZtPtc5* exhibited sequence polymorphism between the parental strains, while *ZtMsr1* is disrupted by a hAT transposon in the 1E4 strain.

### The transcription factor *ZtMsr1* function as hyphal repressor and activator of chlamydospore production, while the protein phosphatase *ZtYvh1* is required for fungal filamentation

Next, we generated mutant lines for the *ZtMsr1, Zt11037, ZtYvh1*, and *ZtPtc5* genes (Fig. S1) and tested their response to heat stress (27°C) (Figs. 3, 4, and S2). Blastospores from 1E4 switched to hyphal growth, while 1A5 produced swollen cells and mature chlamydospores at 48 and 72 hours after incubation (hai), respectively. The ectopic insertion of the 1A5 *ZtMsr1* allele into the 1E4, a strain with *ZtMsr1* disrupted by a TE, induced chlamydospore formation (Fig. 3A). After 24 hai, the 1E4-Ect_*ZtMsr1*-1A5_ mutants displayed a reduction in filamentation and hyphal branches, and an increase in chlamydospore formation, a phenotype that was also observed at later time points. Similarly, deleting the functional *ZtMsr1* in 1A5 resulted in a switch to hyphal growth at 24 hai. Hence, *ZtMsr1* appear to function as a repressor of hyphal growth and activator of chlamydospore production during heat stress (Fig. 3A).

**FIG 3.**
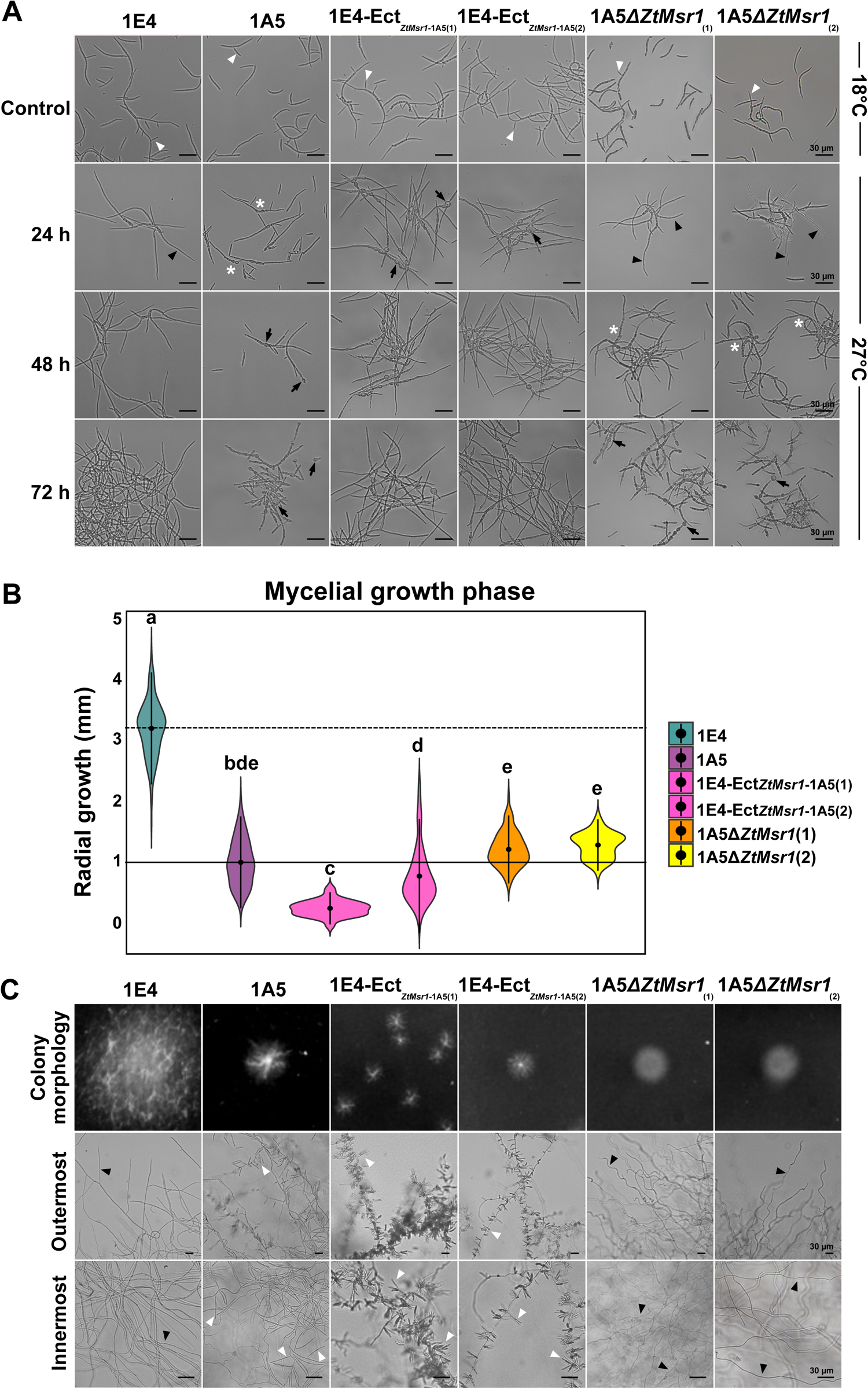
*ZtMsr1* plays a major role in the differential heat stress response observed between the parental strains. (A) Blastospores incubated on nutrient-rich YSB medium at 18°C (control environment) multiply by budding as blastospores (white triangles). The effect of heat stress on *Z. tritici* morphology was observed by exposing the strains at 27°C for 72 h. The 1E4 strain responds to a high temperature by switching mainly to filamentous growth (black triangles), while 1A5 produces pseudohyphae (white asterisks) and chlamydospores (black arrows) as heat stress responses. The 1E4-Ect_*ZtMsr1*-1A5_ mutants were faster to respond to the new environment by producing chlamydospores (black arrows) after 24 h, while the deletion of *ZtMsr1* in the 1A5 strain derepressed filamentation (black triangles). (B) Blastospores incubated on nutrient-limited WA at 18°C switch to hyphal growth. Violin plot demonstrates the radial growth (mm) of the tested strains. The ectopic integration of 1A5-*ZtMsr1* into 1E4 produced smaller colonies compared to 1E4 wild-type. Contrary, the deletion of *ZtMsr1* in the 1A5 did not affect colony size. The dashed or solid horizontal lines indicate the median of the radial growth (mm) of 1E4 and 1A5 strains, respectively. Different letters on the top of the bars indicate a significant difference among the tested strains according to Analysis of Variance (ANOVA). (C) Overview images of colony morphology and light microscopies of the outermost and innermost of the colonies. Colonies from 1E4-Ect_*ZtMsr1*-1A5_ mutants showed extensive blastosporulation resembling 1A5 strain. On the other hand, 1A5*ΔZtMsr1* colonies were hyperfilamented similar to 1E4. Black triangles point to hyphae and white triangles demonstrate areas of blastospore formation. Numbers within brackets represent the mutant line tested.

Additionally, we found that the deletion of *ZtYvh1* drastically reduced filamentation in 1E4 and induced formation of pseudohyphae and chlamydospores at 24 and 48 hai, respectively; however, the morphological response of the 1A5*ΔZtYvh1* null mutant did not differ from its 1A5 wild-type (Fig. 4A). This led us to hypothesize that the 1A5 strain may carry a non-functional allele of *ZtYvh1*. To test this hypothesis, we complemented the 1E4*ΔZtYvh1* with either the 1E4- or 1A5-*ZtYvh1* alleles. Hyphal growth was restored in both 1E4*ΔZtYvh1+*Ect_*ZtYvh1*-1E4_ and 1E4*ΔZtYvh1+*Ect_*ZtYvh1*-1A5_ mutants (Fig. 4A), demonstrating that, despite the high sequence polymorphism exhibited by the parental strains, both *ZtYvh1* alleles appear to be functional. We conclude that Yvh1 is involved in the blastospore-to-hyphae transition in *Z. tritici*, but is not responsible for the differential heat stress responses between the parental strains. Finally, no altered phenotypes were observed for 1E4*ΔZt11037* and 1E4*ΔZtPtc5* compared to its 1E4 wild-type (Fig. S2), therefore no further analyses were performed for these mutants.

**FIG 4.**
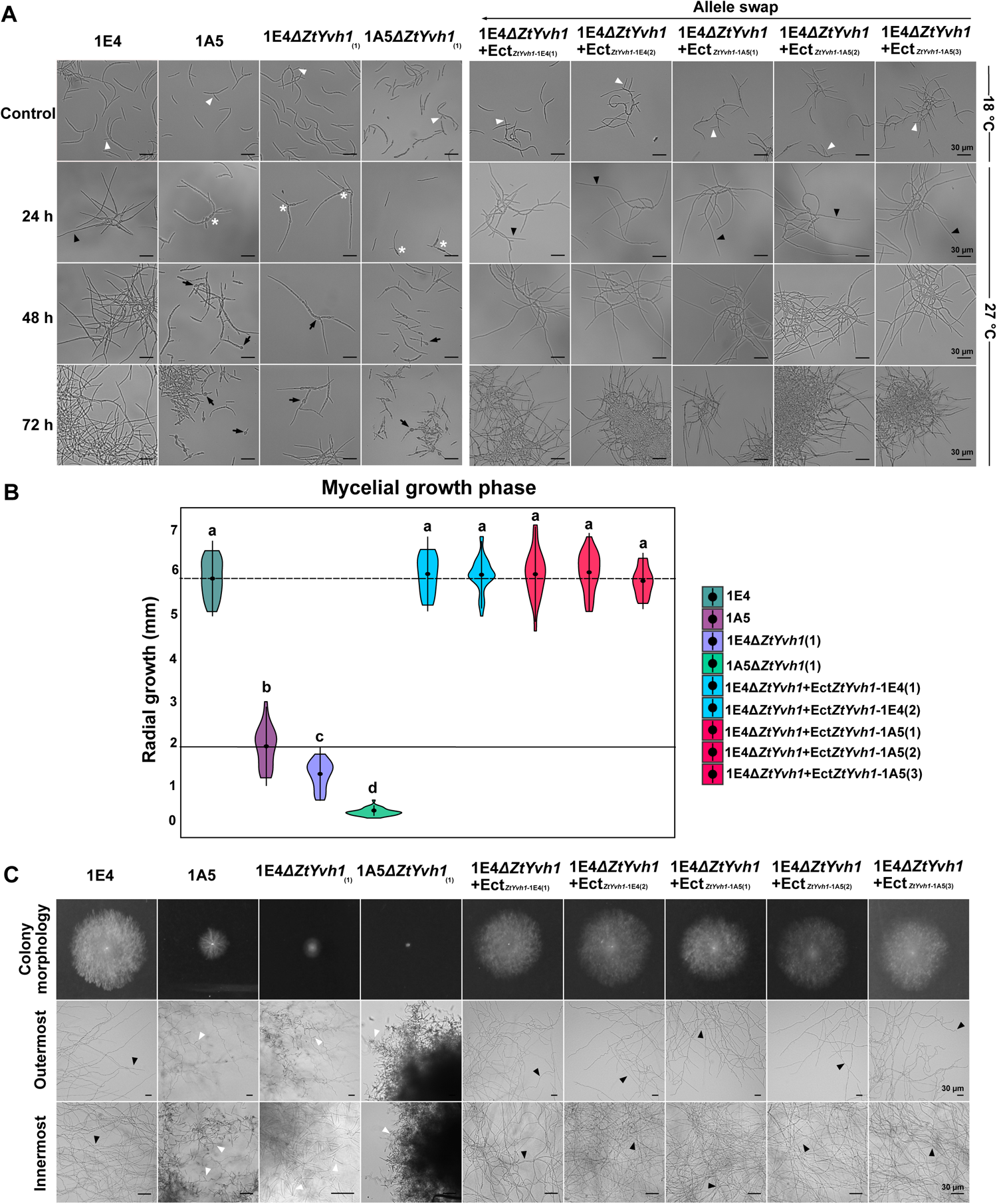
*ZtYvh1* is required for hyphal growth. (A) Blastospores incubated on nutrient-rich YSB medium at 18°C (control environment) multiply by budding as blastospores (white triangles). The effect of heat stress on *Z. tritici* morphology was observed by exposing the strains at 27°C for 72 h. The 1E4 strain undergoes filamentation (black triangles), while 1A5 produces pseudohyphae (white asterisks) and chlamydospores (black arrows) as heat stress responses. Deletion of 1E4-*ZtYvh1* induced a similar response to 1A5, such as pseudohyphae (white asterisks) and chlamydospore (black arrows) formation under high temperatures. However, the 1A5*ΔZtYvh1* null mutant did not differ from its 1A5 wild-type. Complementation of 1E4*ΔZtYvh1* mutant either with 1E4- or 1A5-*ZtYvh1* allele restored the native filamentation (black triangles) of 1E4 strain. (B) Blastospores incubated on nutrient-limited WA at 18°C switch to hyphal growth. Violin plot demonstrates the radial growth (mm) of the tested strains. The deletion of *ZtYvh1* strongly affect colony size independently of the genetic background. The ectopic integration of 1E4- or 1A5-*ZtYvh1* allele into 1E4*ΔZtYvh1* mutant restored the 1E4 mycelial growth. The dashed or solid horizontal lines indicate the median of the radial growth (mm) of 1E4 and 1A5 strains, respectively. Different letters on the top of the bars indicate a significant difference among the tested strains according to Analysis of Variance (ANOVA). (C) Overview images of colony morphology and light microscopies of the outermost and innermost of the colonies. Colonies from 1A5- and 1E4*ΔZtYvh1* mutants showed extensive blastosporulation resembling 1A5 strain. The complemented 1E4-Ect_*ZtYvh1*-1E4_ or 1E4-Ect_*ZtYvh1*-1A5_ mutants restored the 1E4 mycelial growth phenotype. Black triangles point to hyphae and white triangles demonstrate areas of blastospore formation. Numbers within brackets represent the mutant line tested.

### *ZtMsr1* and *ZtYvh1* contribute to vegetative growth and cellular integrity

Because some mutants displayed altered filamentation, we hypothesize that *ZtMsr1* and *ZtYvh1* may also contribute to the mycelial growth phases of *Z. tritici*. To test this hypothesis, we compared the growth of the mutants with their respective wild-types grown on water agar (WA) medium - a nutrient-poor environment that induces hyphal growth (Figs. 3B-C and 4B-C). The vegetative growth inhibition for each tested strain is shown in Table S4 and summarized in Table S5. The 1E4-Ect_*ZtMsr1*-1A5_ mutants showed a growth inhibition of 70% (Figs. 3B and Table S4) and a derepressed blastosporulation on WA compared to the 1E4 wild-type (Fig. 3C). In contrast, deletion of *ZtMsr1* in the 1A5 strain did not affect colony sizes in the nutrient-poor WA environment (Figs. 3B). Although the growth kinetics did not appear to be affected, the 1A5*ΔZtMsr1* produced compact hyperfilamented colonies with repressed blastospore formation (Fig. 3C), a phenotype regularly observed in the 1E4 colonies. This finding confirms that *ZtMsr1* acts as a transcriptional repressor of hyphal growth in *Z. tritici*.

Deletion of *ZtYvh1* significantly impacted vegetative growth of *Z. tritici* independently of the genetic background (Figs. 4B-C). The radial growth of 1E4*ΔZtYvh1* colonies was reduced by 75% compared to 1E4 wild-type on WA plates, while 1A5*ΔZtYvh1* growth was diminished by 82% compared to the 1A5 wild-type strain (Figs. 4B and Table S4). The *ZtYvh1* deletion derepressed blastospore formation in nutrient-poor medium, and colonies of 1E4*ΔZtYvh1* or 1A5*ΔZtYvh1* mutants showed extensive blastosporulation on WA, especially in the innermost part of the colony (Fig. 4C). Nevertheless, the fungal growth suppression observed for the 1E4*ΔZtYvh1* mutant was restored in the 1E4*ΔZtYvh1+*Ect_*ZtYvh1*-1E4_ and 1E4*ΔZtYvh1+*Ect_*ZtYvh1*-1A5_ allele swap mutants (Figs. 4C and Table S4), corroborating that *ZtYvh1* is required for hyphal growth in *Z. tritici*.

Because regulatory circuits controlling heat stress responses are also known to affect cell wall synthesis [8, 44], we next asked if *ZtMsr1* and *ZtYvh1* are also involved in maintaining the cellular integrity of *Z. tritici*. We exposed all tested strains to four different cell wall and membrane stressors (Fig. S3A and Table S5). Deletion of *ZtMsr1* or *ZtYvh1* did not affect the tolerance of *Z. tritici* to hyperosmotic and oxidative stresses caused by sorbitol or H_2_O_2_, respectively (Fig. S3A). Also, the presence/absence of *ZtMsr1* did not influence fungal growth on potato dextrose agar (PDA)-supplemented with SDS, a compound that causes alterations in cell membrane permeability and induces intracellular oxidative stress [45]. In contrast, the presence of *ZtMsr1* does play a crucial role in cell wall sensitivity. The ectopic integration of the 1A5 *ZtMsr1* allele into the 1E4 strain strongly inhibited fungal growth on PDA-supplemented with Congo Red (CR), which inhibits fungal cell wall assembly by binding to chitin and β-1,3 glucan [46]. Interestingly, the 1A5*ΔZtYvh1* null mutant showed substantial tolerance to CR (Fig. S3A). Together with the nature as a transcription factor, these findings suggest that the presence of *ZtMsr1* increases perception of cell wall stresses inhibiting fungal growth on harmful conditions.

In line with previous studies [47, 48], the *ZtYvh1* mutants displayed hypersensitivity to cell-wall perturbing compounds (Fig. S3A). The 1A5*ΔZtYvh1* growth was significantly inhibited on PDA-supplemented with CR, while 1E4*ΔZtYvh1* could not grow in presence of this compound. Both the 1E4*ΔZtYvh1* and 1A5*ΔZtYvh1* mutants showed reduced tolerance to SDS (Fig. S3A). The reconstitution of 1E4*ΔZtYvh1* by the wild-type *ZtYvh1* alleles of either 1E4 or 1A5 restored its growth defect on both CR and SDS (Fig. S3B), confirming that both alleles are functional and the role of Yvh1 for fungal cell wall integrity.

### Deletion of protein kinases from the CWI and HOG pathways result in chlamydospore formation

Due to the role of the MAPK CWI and HOG pathways in thermotolerance and cell wall synthesis in fungi [8, 10, 19], we asked if these two pathways also mediate filamentous growth or chlamydospore production in *Z. tritici* during heat stress. As proof of concept, we tested the previously available IPO323*ΔZtSlt2* and IPO323*ΔZtHog1* MAPK mutants and its wild-type IPO323 strain for their morphological response at 27°C (Fig. S4A). High temperature induced swollen cells and chlamydospore-like structures at 24 hai and 72 hai, respectively, in both IPO323*ΔZtSlt2* and IPO323*ΔZtHog1* mutants. Their morphological responses resembled 1A5, but differed from 1E4 or IPO323 that showed only hyphal growth. Chlamydospores are thick-walled structures that show chitin accumulation in the cell wall [35]. To measure this property, we stained the putative chlamydospore cells with the chitin-binding dye Calcofluor white (CFW). Chitin accumulation was noticeable only in the septa of the suspensor cells (distal part of differentiated hyphae where chlamydospore are formed) or hyphae in the 1E4 and IPO323 strains, whereas the cell wall of the chlamydospore-like cells was highly stained by CFW in the IPO323*ΔZtSlt2* and IPO323*ΔZtHog1*, as observed for chlamydospores produced by 1A5 strain (Fig. S4A), confirming that they were true chlamydospores.

### Chlamydospore formation is a response to the intracellular osmotic stress generated by the heat stress

When exposed to thermal stress, fungal cells rapidly accumulate trehalose to prevent protein denaturation, creating an intracellular hyperosmotic condition [49-51], which induce the efflux and influx of cellular osmolytes in a feedback system [52] governed by cross-talk between the CWI and HOG pathways [10, 18]. We hypothesized that chlamydospore formation is a consequence of a differential fungal perception and/or response to the changing intracellular osmotic environment during heat stress. To test this hypothesis, we inoculated the 1A5, 1E4, and IPO323 strains and their derived mutants on YSB medium amended with 1% sorbitol, used as an osmotic stabilizer, and incubated at 27°C (Fig. 5 and S4B). We found that 1E4 and IPO323 maintained their hyphal growth response, independently of the osmotic condition. In contrast, 1A5 and *ZtMsr1, ZtYvh1*, IPO323*ΔZtSlt2*, and IPO323*ΔZtHog1* mutants that were previously found to undergo chlamydospore formation as the bona-fide heat stress response, switched to filamentous growth in this osmotic stabilized environment, except for IPO323*ΔZtHog1* that recovered its yeast-like growth [53] (Fig. 5 and S4B). Thus, we concluded that the intracellular osmolarity functions as a crucial signal that determines the regulatory cascade controlling the morphogenic responses under heat stress.

**FIG 5.**
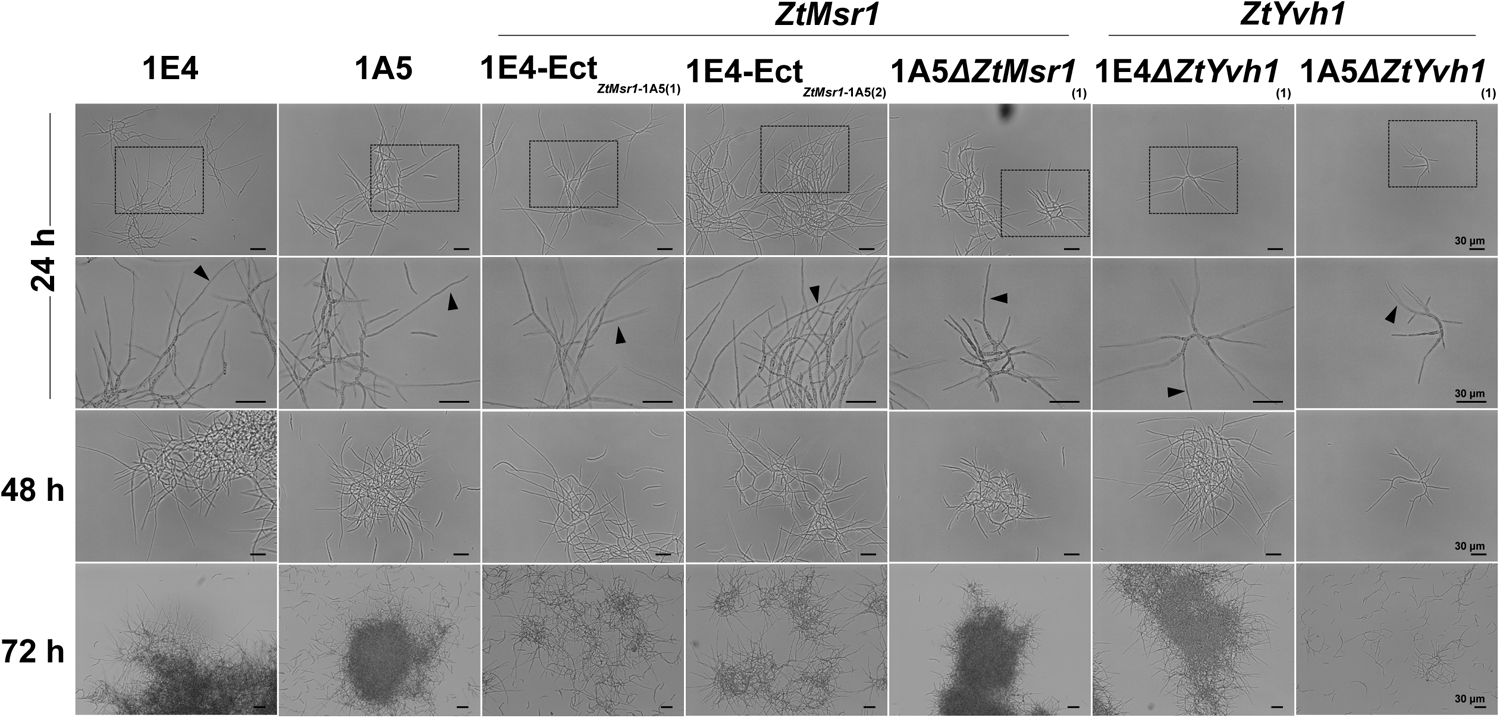
Chlamydospore formation is a morphological response to the intracellular osmotic upshift generated during heat stress. Fungal spores were grown at 27°C for 72h on YSB-supplemented with 1M sorbitol. The ability of 1A5 and all mutant lines producing chlamydospores as a temperature-dependent response was abolished. In this osmotically stabilized condition, the 1A5 strain and the *ZtMsr1* and *ZtYvh1* mutant lines solely produce filamentation (black triangles), as heat stress morphotype.

## DISCUSSION

Global warming is likely to have a significant effect on fungal populations, favoring thermotolerant strains or those producing heat-resistant survival structures. This study extends previous observations on temperature-dependent morphogenesis in fungi [54-56]. Previously, we showed that Swiss *Z. tritici* strains produce chlamydospores as a temperature-dependent morphotype [35]. Here, we demonstrated that *Z. tritici* populations from around the world can grow as hyphae or chlamydospores in response to heat stress. In addition, QTL mapping allowed us to identify genes regulating morphogenetic transitions in *Z. tritici*. Our data proposes a complex mechanism underlying morphogenic transitions in response to temperature stress (Fig. 6).

**FIG 6.**
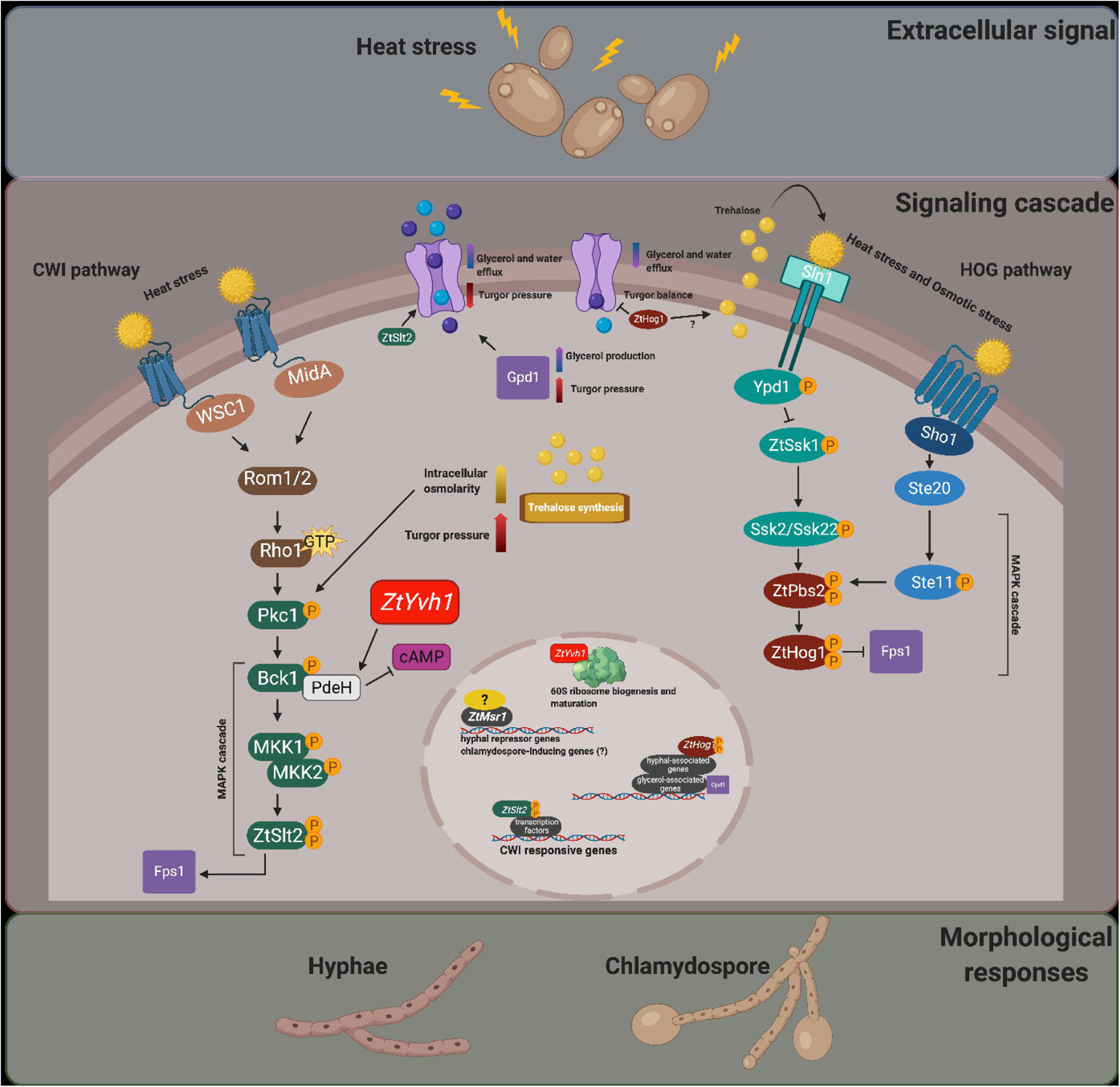
A proposed model for the cross-talk between the CWI and the HOG pathways showing the contribution of *ZtMsr1* and *ZtYvh1* to temperature-dependent morphogenesis in *Zymoseptoria tritici*. Heat stress is known to induce accumulation of cytoplasmic trehalose, causing an increase in intracellular osmolarity and turgor pressure. The MAPK Pck1 from the cell wall integrity (CWI) pathway perceives this signal, which in turn activates downstream MAPK genes by phosphorylation. The phosphorylated-Slt2 is translocated into the nucleus and activates genes responsible for the maintenance of cell integrity. ZtMsr1 and ZtYvh1 are required for morphological transitions in response to heat stress. Yvh1 has been previously demonstrated to activate the phosphodiesterase PdeH, which physically interacts with MAPK BcK1 from CWI and negatively regulates cAMP. Yvh1 also participates in the 60S ribosome biogenesis and maturation and thereby may contribute to the decoding of messenger RNAs affecting cellular processes in fungi, including vegetative growth. *Hog1* is activated via the Sho1-dependent pathway during heat stress. *ZtHog1* has also been demonstrated to be essential to activate hyphal-associated genes in *Z. tritici*. It has been proposed that *Hog1* contributes to the extracellular translocation of trehalose, although this mechanism is not yet understood. The efflux of trehalose is then perceived by Sln1, the second branch of the HOG pathway. The signaling through Sln1 activates Hog1, which in turn activates Gpd1 responsible for glycerol biosynthesis. The intracellular accumulation of glycerol also increases turgor pressure. To counterbalance the intracellular pressure, Slt2 phosphorylates Fps1, which opens the aquaporin water/glycerol channel to release glycerol and water. Once the normal osmotic conditions are reestablished, Hog1 phosphorylates Fps1 to close the channel. As demonstrated in this study, chlamydospore formation is suppressed in an osmotic stabilized environment, indicating that *Z. tritici* strains may perceive differently the intracellular osmolarity, which in turn regulates the signaling cascade in a different manner resulting in hyphal or chlamydospore growth. Image created with BioRender.com.

The ability to switch among different cell morphologies is critical for the survival of most filamentous fungi, with different types of cells playing different roles during their life cycles. For instance, hyphal growth is essential for virulence in *Z. tritici* [57], and a mutant unable to transition to the hyphal morphology is non-pathogenic [53]. Chlamydospores are survival structures that allow fungi to persist as resting spores during harsh conditions and to infect host tissues once the environment becomes more conducive to growth [35, 58, 59]. The different cellular morphologies observed in *Z. tritici* worldwide field populations, including blastospores, hyphae, and chlamydospores, may differentially affect the fitness of *Z. tritici* strains at different points in their life cycle, with the ability to switch among morphologies governed by natural genetic variations. We showed that these variations permit the flexibility and survival of a fungal population facing environmental changes [35]. Therefore, these findings motivated us to investigate the genetic basis of temperature-dependent morphogenesis in this fungus.

We identified a single QTL on chromosome 12 containing only eight genes that explained 25% of the overall variation for a temperature-dependent morphological switch. This illustrates that morphological changes can be inherited as quantitative traits in *Z. tritici* [32]. Two genes in this QTL, a novel transcription factor named *ZtMsr1* and a protein phosphatase named *ZtYvh1*, regulate blastospore-to-hyphae/chlamydospore transition in response to heat stress in *Z. tritici*. Zinc cluster transcription factors (TFs) like *ZtMsr1* are key players in the signal transduction pathways regulating a plethora of cellular and stress responses in fungi [60, 61]. In this study, the deletion of *ZtMsr1* in the 1A5 strain activated hyphal growth, whereas the insertion of the 1A5-*ZtMsr1* allele into the 1E4 strain (in which *ZtMsr1* is naturally disrupted by a TE), induced chlamydospore formation. Furthermore, after incubation onto a mycelial-inducing environment, the 1E4-Ect_*ZtMsr1*-1A5_ mutants showed impaired hyphal growth and derepressed blastosporulation like 1A5, while 1A5*ΔZtMsr1* mutants exhibited hyperfilamented colonies similar to 1E4. Taken together our findings underpin that *ZtMsr1* is positively required for chlamydospore formation and hyphal repression in *Z. tritici*. Until now, *Zt107320* was the only zinc cluster TF described to regulate the blastospore-to-hyphae transition in *Z. tritici*; however, that response was to different sources of carbon [62]. Similar to other dimorphic fungi, the zinc cluster proteins may act as activators, repressors, or as both activators and repressors for specific genes. For example, *Stb5* acts as an activator and a repressor in the budding yeast *Saccharomyces cerevisiae* in response to oxidative stress [63]. The complete repression of hyphal growth or the sole induction of chlamydospore formation in *ZtMsr1* mutants likely depends on contributions from other genetic modifiers, consistent with the polygenic nature of morphological transitions in fungi. Additional studies will be needed to identify *ZtMsr1*-regulated genes or determine if *ZtMsr1* can be self-regulated or regulate other genes within the chromosome 12 QTL.

Contrary to *ZtMsr1*, the protein phosphatase *ZtYvh1* orthologs are well-characterized in eukaryotes. Deletion of *Yvh1* in different fungi causes defects for a multitude of fungal development processes, including vegetative growth and cellular integrity [38-40, 47, 48]. Nevertheless, this is the first time that an *Yvh1* ortholog has been shown to regulate morphological stress responses. Deletion of the 1E4 *ZtYvh1* allele significantly repressed filamentation and induced chlamydospore formation during heat stress, whereas deletion of *ZtYvh1* did not affect the morphological responses of 1A5. At first, this result suggested that the 1A5 *ZtYvh1* phosphatase was not functional. However, the reconstitution of the 1E4*ΔZtYvh1* mutant with either the 1E4- or 1A5-*ZtYvh1* alleles restored the WT filamentation phenotype and abolished chlamydospore formation upon heat stress, demonstrating that ZtYvh1 is involved in the dimorphic switch, but is not responsible for the differential heat stress responses between the parental strains. Consistent with the pleiotropic role of this phosphatase, the deletion of *ZtYvh1* also affected the vegetative growth of *Z. tritici*, independently of the genetic background. Since the deletion of *ZtYvh1* orthologs in *C. albicans CaYvh1, S. cerevisiae ScYvh1*, and *M. oryzae MoYvh1* also cause defects in fungal growth [39, 40, 48, 64], we believe that the role of *ZtYvh1* does not differ among these fungi. *ScYvh1* was demonstrated to be involved in 60S ribosomal subunit biogenesis, where it is recruited to the pre-60S to facilitate the release of the Mrt4 assembly factor from the ribosome stalk of the maturing 60S particles [65-67]. Genes involved in ribosome biogenesis are regulated in response to environmental stresses [68]. *ScYvh1* mutants exhibit defective 60S ribosome biogenesis because the signal transduction delayed translation-competent 60S subunit affects the folding of nascent polypeptides and decoding of messenger RNA [47, 65, 66]. We speculate that a similar mechanism leads to the impaired vegetative growth observed for the *ZtYvh1* mutants, especially during a temperature shift [38, 39, 69].

Another phenotype typically observed in *Yvh1* mutants is the hypersensitivity to cell-perturbing agents [47, 48]. The regulation of Bck1-Mkk1-Slt2 MAPK cascade is thought to be crucial for CWI. In *M. oryzae, MoYvh1* functions in the maintenance of CWI via the regulation of the MAPK pathway [48]. Consistent with the findings in *ΔMoYvh1*, the *ΔZtYvh1* null mutants were sensitive to both CR and SDS, suggesting that ZtYvh1 might be required for cell wall integrity in *Z. tritici*. Additionally, we pinpointed that ZtMsr1 increases fungal perception to cell wall stresses, for example, the cell damage caused by the increased temperature. While the 1E4-Ect_*ZtMsr1*-1A5_ mutants were hampered by the CWI inhibitor CR, the deletion of *ZtMsr1* caused tolerance to this compound. Taken together all the above-mentioned results, we suspected that the activation of CWI signaling pathway may allow the fungus to sense and response to cell wall stresses that arise during temperature upshift.

Upon heat stress, fungal cells accumulate trehalose, which protects native proteins against thermal-denaturation [70, 71], but also increases the intracellular osmolarity [11, 72, 73]. The cytoplasmic accumulation of this osmolyte compound causes hypotonic stress and stretches the plasma membrane, a condition sensed by Pck1 MAPK which in turn activates the CWI pathway [49, 74]. Preventing trehalose synthesis diminishes CWI signaling during heat stress [49]. In parallel, *Hog1* is rapidly phosphorylated through the Sho1-Pbs2-Hog1 MAPK cascade upon a temperature shift [18, 19]. Although the details of the cross-talk between the HOG and CWI pathways are not understood, mutants deleted in components of the CWI pathway are hypersensitive to heat stress. The deletion of *ZtSlt2* is reported to promote large and swollen cells in filamentous fungi and oomycetes, including in *Z. tritici* [25-28]. We showed that these “swollen hyphae” are, in fact, chlamydospores. An earlier QTL study of temperature sensitivity in *Z. tritici* identified the Pbs2 MAPK [32], suggesting that the HOG pathway may also play a role in thermal adaptation in this fungus. The deletion of *ZtHog1* was earlier demonstrated to abolish the yeast-to-hyphae transition in *Z. tritici* [53]. Here, we showed that deletion of *ZtHog1* mutant induced chlamydospore formation, which supports our findings that the filament-specific components need to be inactivated for the formation of chlamydospores, as demonstrated for *ZtMsr1* and *ZtYvh1* mutants. The deletion of *Hog1* also causes osmosensitivity in different yeasts and filamentous fungi [75-78], including in *Z. tritici* [53]. Osmotic stabilization of the plasma membrane in the presence of a defective cell wall can prevent stimulation of the CWI pathway, and this would avert the regulatory cascade leading to hyphal or chlamydospore formation. Using an osmotic stabilized environment, we found that all the strains producing chlamydospores, including 1A5, underwent filamentation, except for the IPO323*ΔZtHog1* mutant. Surprisingly, this MAPK mutant recovered its yeast-like growth in the osmotic stabilized condition, demonstrating that the intracellular osmotic stress is, in fact, the signal inducing chlamydospore formation thought the stimulus of the CWI pathway; however, the induction of filamentation requires the integrity of HOG pathways in *Z. tritici*. This finding leads us to hypothesize that chlamydospores might be induced under any conditions that cause an osmotic upshift during the fungal lifecycle, in line with our previous discovery that chlamydospores are produced during the disease cycle of *Z. tritici* [35]. In summary, we concluded that *ZtMsr1* increases perception of cell wall stresses inhibiting fungal hyphal on harmful condition, such when temperature rises – a condition that also cause fungal cell wall damage. In turn, when cell wall integrity is compromised, *ZtMsr1* might be activated and regulate the chlamydospore-inducing genes as a stress response survival strategy.

Our data illustrate the stunning complexity of mechanisms affecting temperature-dependent morphogenesis, including the effects of naturally occurring genetic variants involved in the cross-regulation of MAPK signaling cascades. The results presented here provide a solid basis for future studies on the intracellular processes driving morphological transitions in fungi. Despite some remaining questions, the general implication of our findings is clear: *ZtMsr1* and *ZtYvh1* contribute to temperature-dependent responses in the fungal plant pathogen *Zymoseptoria tritici*.

## MATERIAL AND METHODS

### Fungal isolates and growth conditions

A total of 372 isolates of *Z. tritici* were used to assess the morphological stress response in this fungus (Tables S1 and S2). The two Swiss *Z. tritici* strains ST99CH_1A5 (abbreviated as 1A5) and ST99CH_1E4 (abbreviated as 1E4), and mutant lines derived from these strains were also used here (Fig. S1). The knocked-out IPO323*ΔZtSlt2* [27] and IPO323*ΔZtHog1* [53] mutants were provided by Marc-Henri Lebrun (National Institute of Agricultural Research – INRA). Because the MAPK mutants were generated in the genetic background of IPO323 [79], this strain was also used as a control. Fungal cells were routinely retrieved from glycerol at - 80°C and grown on YSB (10 g/L yeast extract, 10 g/L sucrose, and 50 μg/mL kanamycin sulfate; pH 6.8) medium at 18 °C for four days. Cell concentrations were determined by counting blastospores using the KOVA cell chamber system (KOVA International Inc., USA) and kept on ice until required for the phenotypic assays.

### Phenotyping for the temperature-dependent morphotypes

Blastospore suspensions of each tested isolate were added to a final concentration of 10^5^ blastospores/mL on YSB and incubated at 27°C. After 72 hours of incubation, an aliquot was taken and checked by light microscopy using a Leica DM2500 microscope with LAS version 4.6.0 software. Isolates were scored for their stress response to grow as chlamydospores (score=0), hyphae (score=1), or as a mixture of chlamydospores and hyphae (score=2) under the heat stress (Tables S1 and S2).

To determine the global distribution of morphological stress responses, we analyzed 141 isolates of *Z. tritici* collected from single wheat fields between 1990 and 2001 in four distinct locations: Australia, Israel, Switzerland, and Oregon (USA) [80]. In Oregon, isolates were sampled from the resistant cultivar Madsen (Oregon R) and the susceptible cultivar Stephens (Oregon S).

To analyze the genetic architecture of temperature-dependent morphogenesis, we used a *Z. tritici* mapping progeny population consisting of 261 offspring individuals from the cross between 1A5 and 1E4 [81]. These two parental strains were sampled from the same naturally infected wheat field in Switzerland in 1999 [82] and differ for several traits, including their morphological stress response [35].

### Genotype data and QTL mapping

SNP data of 261 offspring isolates from the cross between 1A5 and 1E4 were obtained from RAD sequencing data [81]. The genetic maps generated from the SNPs segregating in the progeny population were assembled using the 1A5 reference genome [83, 84]. SNP markers were filtered to include only SNPs called among the 1A5 and 1E4 parental genomes. This provided 35’030 SNP markers in the 1A5 × 1E4 cross with an average marker distance of 1’145 bp (equivalent to 0.31 cM). The QTL analysis was based on the phenotypic score given to each offspring isolate, as demonstrated in Tables S2 and S6. The Single-QTL genome scan using the standard interval mapping (SIM) was performed in the R/qtl package [85] to improve the marker regression method by estimating pseudomarkers in between true markers. The significance thresholds of logarithm odds (LOD) of the QTLs were based on 1000 permutation tests across the entire genome, followed by Bayesian credible intervals used to calculate 95% confidence intervals of the QTL. Genes within the 95% confidence interval were identified according to the genome annotations of the reference parental strains [84].

### Identification of candidate genes within the QTL confidence interval

All genes within the QTL confidence interval had their genome sequences compared to identify variants between the 1A5 and 1E4 parental strains using AliView software [86]. We evaluated genes for the presence of non-synonymous SNPs and other sequence variation either in the protein-encoding sequence or the 5’- or 3’-UTR. The candidate genes were BLASTed to the NCBI database (https://blast.ncbi.nlm.nih.gov/Blast.cgi) to confirm their functional domains and to search for orthologs in other fungal species. Synteny of the 1A5 and 1E4 genome sequences in the QTL region was analyzed using pairwise blastn on repeat-masked genomic sequences and visualized using the genoPlotR package in R [87].

### Plasmid constructions and transformations

DNA assemblies were conducted with the In-Fusion HD Cloning Kit (Takara BIO) following the manufacturer’s instructions. A summary of the plasmids and their constructions is given in Fig. S1 and primers used in this study are listed in Table S7. *Z. tritici* cells were transformed via *Agrobacterium*-mediated transformation according to the protocol adapted by Meile et al., 2017 [88]. We confirmed the mutant lines by a PCR-based approach using a forward primer specific to the upstream sequence of the inserted cassette and a reverse primer specific to the selective gene. We determined the copy number of the transgene by quantitative PCR (qPCR) on genomic DNA extracted with the DNeasy Plant Mini Kit (Qiagen). Lines with a single insertion were selected for further experiments.

### Phenotypic characterization of the mutants

Blastospore suspensions of each strain were inoculated into three flasks containing YSB at a final concentration of 10^5^ blastospores/mL and incubated at 18°C (as control) or 27°C for three days. The cell morphology was observed by light microscopy every 24 hours after incubation (hai) until 72 hai. We used the morphology of the 1A5 or 1E4 strains as references.

To test for altered vegetative growth, we used water agar (WA – 12 g/L agar, and 50 μg/mL kanamycin sulfate) medium to induce hyphal growth. 200 μL of a blastospore suspension of each tested strain was plated at a final concentration of 2×10^2^ blastospores/mL on five independent WA plates and incubated in the dark at 18°C for 15 days post-inoculation (dpi). Because mycelial growth on WA plates exhibited poor color contrast, the colony diameters were measured manually. All measurements included at least 20 colonies. The colony diameter values were divided by two to generate the radial growth (mm) values, which were plotted in a violin plot ggplot2 package from R [89]. Analysis of variance (ANOVA) was performed using the agricolae package [90]. The radial growth (mm) of 1A5 and 1E4 strains was used to calculate the percentage of growth inhibition (% of reduction of growth radius) of each the mutant. A t-test statistic was used to test the hypothesis that mutants were affected in their vegetative growth compared to the wild-type strains.

To assess the contribution of the candidate genes to the cell wall integrity, we exposed blastospores of the tested strains to different stress conditions, including oxidative stress (1 mM of hydrogen peroxide – H_2_O_2_), osmotic stress (1M sorbitol), cell wall stress (2 mg/mL Congo red), and plasma membrane stress (0.01% sodium dodecyl sulfate). Blastospore suspensions of each strain were serially diluted to 4×10^7^, 4×10^6^, 4×10^5^, and 4×10^4^ blastospores/mL and drops of 3.5 μL were plated onto five independent potato dextrose agar plates (39 g/L potato dextrose agar, and 50 μg/mL kanamycin sulfate) amended with the above-mentioned stresses, and incubated at 18°C in a dark room. Blastospore suspensions plated only onto PDA plates were used as a control. Colony phenotypes were assessed in a qualitative manner using digital images taken at 6 dpi.

### Osmotic stabilization

Blastospore suspensions of the mutant lines and their respective wild-type strains were inoculated onto YSB medium amended with 1M sorbitol at a final concentration of 10^5^ blastospores/mL and incubated at 27°C. Flasks containing only YSB medium were used as control. An aliquot was taken from each flask at 24, 48, and 72 hai to monitor cell morphology by light microscopy.

### The use of MAPK mutants as proof of concept

To evaluate the role of the MAPK CWI and HOG pathways in thermotolerance [8, 10, 19], we tested whether these two MAPK pathways also play a role in the temperature-dependent morphogenesis described here using IPO323*ΔZtSlt2* and IPO323*ΔZtHog1* mutants, as proof of concept. Blastospore suspensions of each strain were inoculated onto YSB at a final concentration of 10^5^ blastospores/mL and incubated at 18°C (as control) or 27°C. The 1A5, 1E4 and IPO323 *Z. tritici* strains were used as references. Cell morphologies were analyzed by light at 24, 48, and 72 hai. Cells harvested at 72 hai were also fixed with 70% (v/v) ethanol for 30 min, followed by three washes with phosphate-buffered saline (PBS). The cells were stained with 1 μg/mL of chitin-binding dye Calcofluor white (CFW) (Sigma-Aldrich Chemie Gmbh, Munich, Germany) for 15 min. The stained cells were viewed with a Leica DM2500 fluorescence microscope using a UV filter system for CFW consisting of a BP excitation filter at 340-380 nm and a long pass emission filter (>425 mm).

### Data Availability

The genotypic and phenotypic data and marker information of the 1A5 × 1E4 cross-population are available in Table S6.

## ACKNOWLEDGEMENTS

This study was financed in part by the Coordenação de Aperfeiçoamento de Pessoal de Nível Superior – Brasil (CAPES) – Finance Code 001. We acknowledge Marc-Henri Lebrun from the French National Institute of Agricultural Research (INRA) for kindly providing the MAPK mutants used in this study. We thank Michael Habig and Jason Rudd for the critical reading of the manuscript. We also thank Julien P. L. Alassimone, Dominik Vetsch, and Anja Kunz for their assistance with the experiments.

## SUPPLEMENTAL MATERIAL SUPPLEMENTAL FIGURES

**FIG S1. Mutants generated in this study**. (A) Representation of the genetic composition of the 1A5 and 1E4 parental strains. (B-I) The schematic diagram for each mutant line is represented by the wild-type genetic background used for the *Agrobacterium*-mediated transformation (first line), the plasmid construction (second line), and the transformant (third line). To create the construct for ectopic integration of *ZtMsr1*_1A5_ into the 1E4 genetic background, a fragment containing *ZtMsr1*_1A5_, 1 Kb upstream of the start codon, and 1 Kb downstream of the stop codon were amplified from 1A5 genomic DNA. The pES1 plasmid (obtained from E. H. Stukenbrock, Kiel University, unpublished) carrying the hygromycin resistance cassette and used as a selectable marker was linearized with *Xba*I and *Him*III (New England Biolabs). The two fragments were assembled into pES1 resulting in pES1-Ect_*ZtMsr1*-1A5_. To knock-out the genes *Zt11059, ZtYvh1*, or *ZtPtc5* in the 1E4 strain, 1 Kb of both flanking regions for each gene were amplified from the 1E4 genomic DNA. The pES1 plasmid was digested with *Kpn*I and *Sbf*I (New England Biolabs) for plasmid linearization, and three fragments of each construct were assembled into pES1 carrying the hygromycin resistance cassette and used as selectable marker, resulting in pES1-1E4*ΔZt11059*, pES1-1E4*ΔZtYvh1*, and pES1-1E4*ΔZtPtc5*. The *ZtMsr1* and *ZtYvh1* genes were also knocked-out in the 1A5 strain. The linearized pES1 plasmid with *Kpn*I and *Sbf*I (New England Biolabs) and the two flanking regions of each gene were assembled into pES1, resulting in pES1-1A5*ΔZtMsr1* and pES1-1A5*ΔZtYvh1*. To create the constructs for ectopic integration of the *ZtYvh1*_1E4_ or *ZtYvh1*_1A5_ genes into the 1E4*ΔZtYvh1* mutant strain, the pCGEN plasmid carrying the geneticin resistance cassette as a selective marker [33] was linearized with *Kpn*I (New England Biolabs). A fragment containing the *ZtYvh1*_1E4_ or *ZtYvh1*_1A5_ genes and 1 Kb of their respective flanking regions were amplified from 1E4 or 1A5 genomic DNAs, respectively, and cloned into pCGEN, resulting in pCGEN-1E4-Ect_*ZtYvh1*-1E4_ or pCGEN-1E4-Ect_*ZtYvh1*-1A5_. We confirmed the mutant lines by a PCR-based approach using a forward primer specific to the upstream sequence of the inserted cassette and a reverse primer specific to the selective gene. All primers used for plasmid constructions and validations are indicated by the arrows and their sequences are described in Table S7.

**FIG S2. *Zt11073* and *ZtPtc5* do not contribute to temperature-dependent morphogenesis in *Zymoseptoria tritici***. (A) Blastospores incubated on nutrient-rich YSB medium at 18°C (control environment) or 27°C (heat stress) for 72h. Both knocked-out *Zt11073* and *ZtPtc5* mutants did not differ in their morphological response compared to its 1E4 wild-type. White and black triangles point to blastospores and filamentous hyphae, respectively. White asterisks and black arrows indicate pseudohyphae and chlamydospores formed by the 1A5 strain in response to the heat stress. Numbers within brackets represent the mutant line tested.

**FIG S3. Genetic contributions of *ZtMsr1* and *ZtYvh1* to the cellular integrity**. (A) Blastospores were plated on PDA supplemented with cell wall or plasma membrane stressors and grown at 18°C for six days. Deletion of *ZtMsr1* or *ZtYvh1* did not affect the tolerance of *Z. tritici* to sorbitol or hydrogen peroxide (H_2_O_2_). The ectopic integration of 1A5-*ZtMsr1* into 1E4 promoted hypersensitivity to Congo Red (CR). Deletion of *ZtYvh1* affected fungal tolerance to cell wall stressors. 1E4*ΔZtYvh1* growth was suppressed in the presence of CR, and both 1E4*ΔZtYvh1* and 1A5*ΔZtYvh1* mutants were inhibited in the presence of Sodium Dodecyl Sulfate (SDS). (B) Reconstitution of the 1E4*ΔZtYvh1* mutant either with 1E4- or 1A5-*ZtYvh1* allele restored the growth defect in presence of CR and SDS compounds. Numbers within brackets represent the mutant line tested.

**FIG S4. Deletion of genetic components of the MAPK CWI and HOG pathways derepressed chlamydospore formation during heat stress**. (A) Strains were incubated at 18°C (control) or 27°C (heat stress). High temperature induced swollen cells and chlamydospore-like structures in both IPO323*ΔZtSlt2* and IPO323*ΔZtHog1* mutants. Their morphological responses resembled 1A5, but differed from 1E4 or IPO323 that showed only hyphal growth, as thermal response. Cell walls were stained with the chitin-binding dye Calcofluor White (CFW) (blue color). Hyphae, pseudohyphae, or suspensor cells (differentiated hyphae where chlamydospores are usually attached) showed a weaker fluorescence for CFW compared to chlamydospores produced by 1A5, IPO323*ΔZtSlt2*, and IPO323*ΔZtHog1* strains. White and black triangles indicate blastospores and hyphal growth, respectively. White asterisks indicate pseudohyphae, while black arrows point to chlamydospore cells. (B) Fungal spores grown at 27°C for 72 h on YSB-supplemented with 1M sorbitol. The ability of 1A5 and IPO323*ΔZtSlt2* and IPO323*ΔZtHog1* mutants to produce chlamydospores as a temperature-dependent response was abolished in this osmotically stabilized condition.

## SUPPLEMENTAL TABLES

**Table S1**. Phenotypic data of 141 *Zymoseptoria tritici* isolates from five global populations.

**Table S2**. Phenotypic data of 231 progeny isolates from the cross between 1A5 and 1E4 Swiss *Zymoseptoria tritici* strains used for quantitative trait locus (QTL) mapping analysis.

**Table S3**. Genes identified within the QTL confidence interval on chromosome 12 for the differential morphological stress responses in the 1A5 × 1E4 cross.

**Table S4**. Effect of the candidate genes on the mycelial growth phase of *Zymoseptoria tritici* and the percentage of growth inhibition on nutrient-poor condition.

**Table S5**. Summary of the genetic contribution of *ZtMsr1* and *ZtYvh1* to the morphological stress response and cell integrity of *Zymoseptoria tritici*.

**Table S6**. Genotypic and phenotypic data and marker information of the 1E4 × 1A5 cross-population.

**Table S7**. List of primers used in this study.

