## Supplementary figures and images for "A transcription factor and a phosphatase regulate temperature-dependent morphogenesis in a fungal plant pathogen"

### Fig. S1

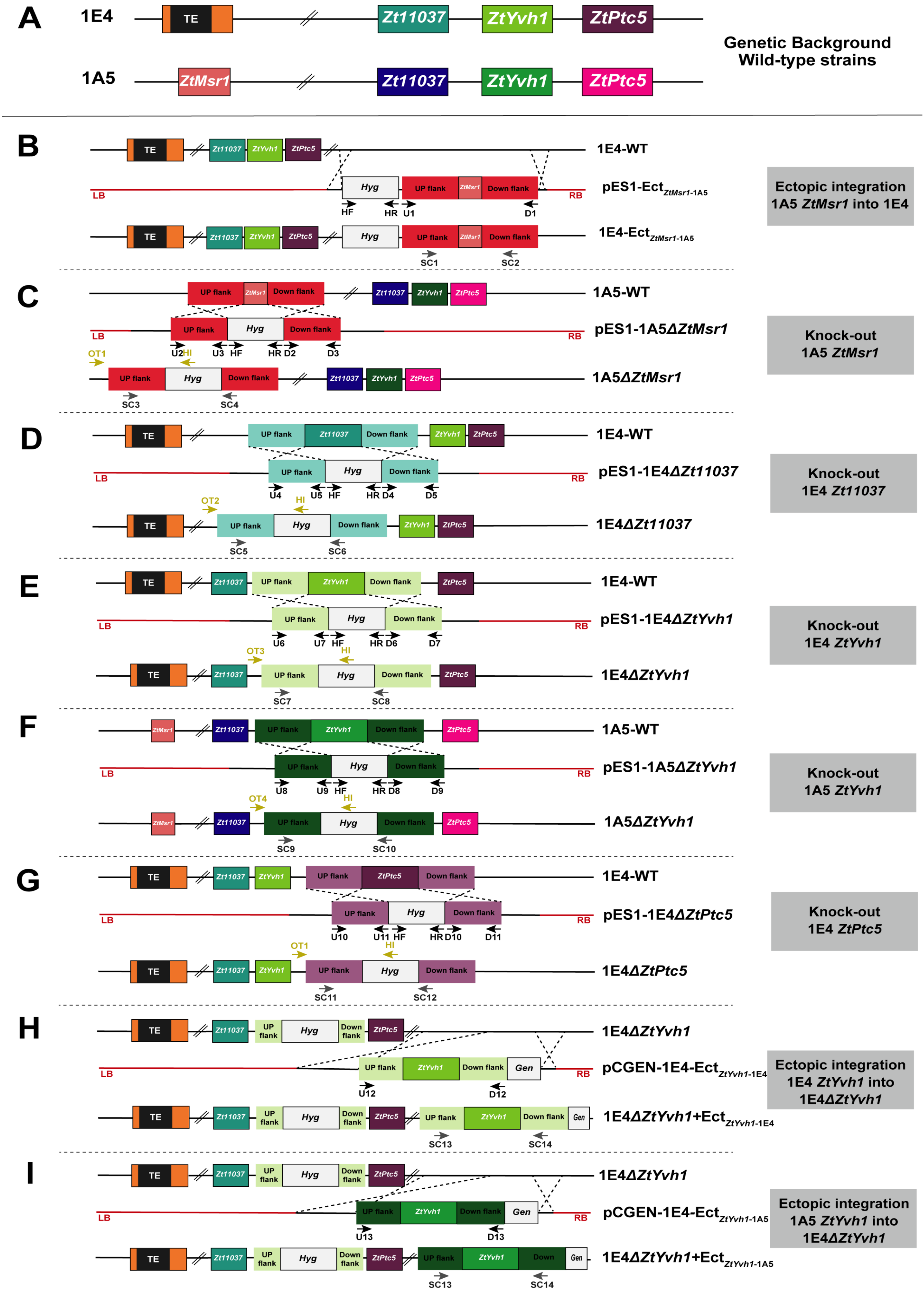

### Fig. S2

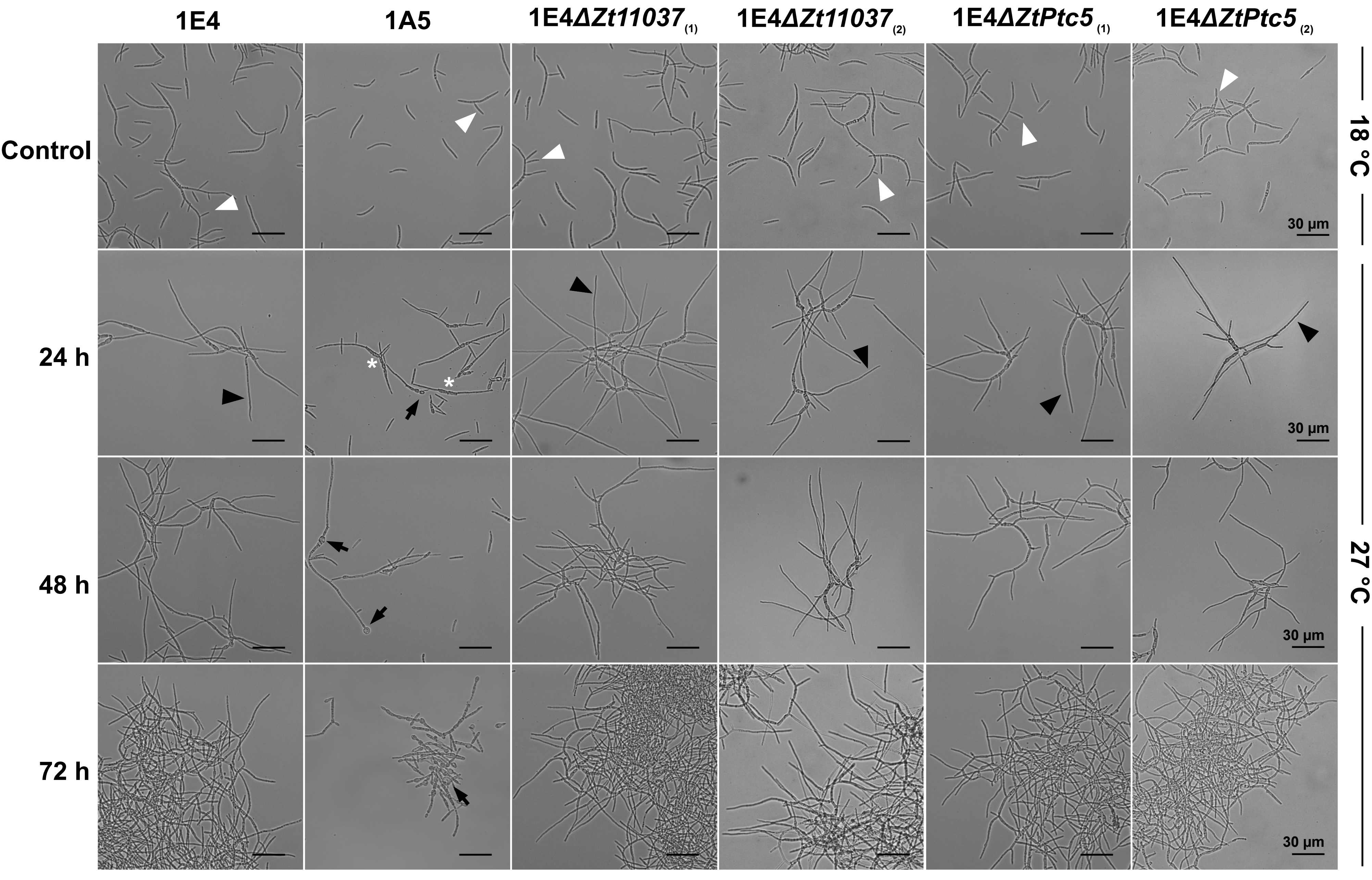

### Fig. S3

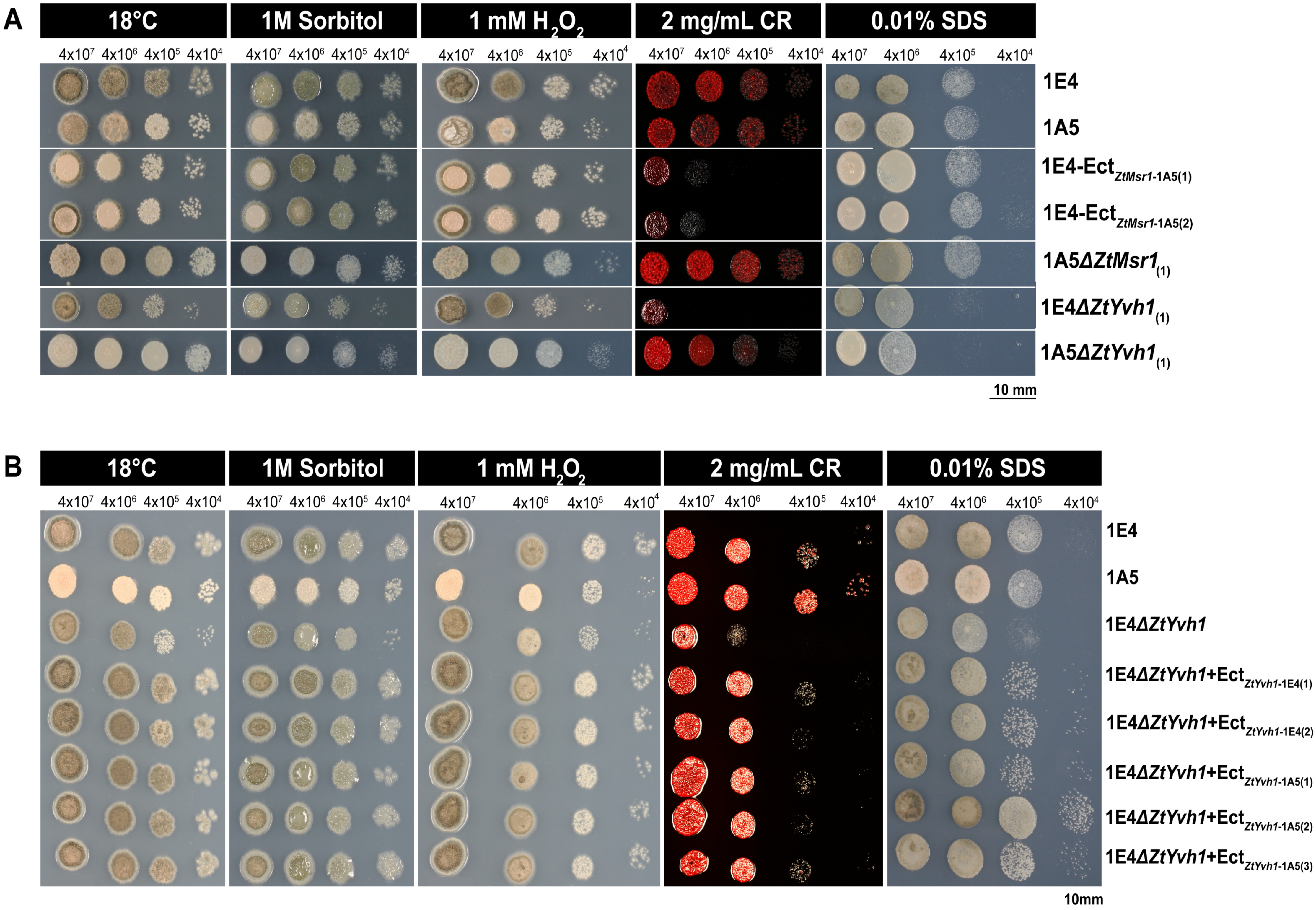

### Fig. S4

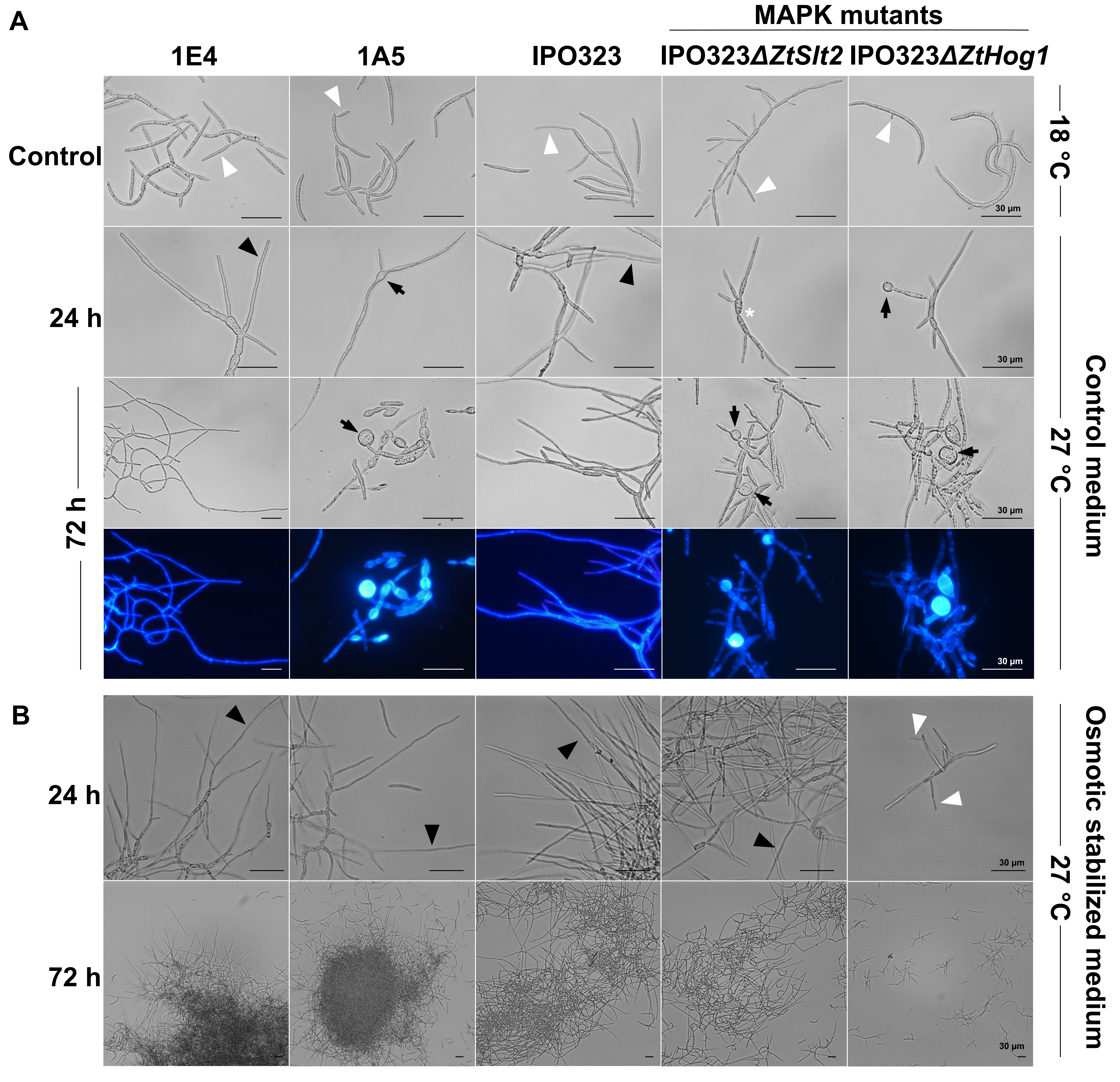
